# A physiologically inspired hybrid CPG/Reflex controller for cycling simulations that generalizes to walking

**DOI:** 10.1101/2024.09.06.611594

**Authors:** Giacomo Severini, David Muñoz

**Author notes:** Corresponding Author: Giacomo Severini.

## Abstract

Predictive simulations based on explicit, physiologically inspired, control policies, can be used to test theories on motor control and to evaluate the effect of interventions on the different components of control. Several control architectures have been proposed for simulating locomotor tasks, based on fully feedback, reflex-based, controllers, or on feedforward architectures mimicking the Central Pattern Generators. Recently, hybrid architectures integrating both feedback and feedforward components have been shown to represent a viable alternative to fully feedback or feedforward controllers. Current literature on controller-based simulations, however, almost exclusively presents task-specific controllers that do not generalize across different tasks. The task-specificity of current controllers limits the generalizability of the neurophysiological principles behind such controllers. Here we propose a hybrid controller for predictive simulations of cycling based where the feedforward component is based on a well-known theoretical model, the Unit Burst Generation model, and the feedback component includes a limited set of reflex pathways, expected to be active during submaximal steady cycling. We show that this controller can simulate physiological cycling patterns at different desired speeds. We also show that the controller can generalize to walking behaviors by just adding an additional control component for accounting balance needs. The controller here proposed, although simple in design, represent an instance of physiologically inspired generalizable controller for cyclical lower limb tasks.

**Author Summary:** Predictive simulations allow to synthesize human movements and their associated biomechanical quantities without using experimental data. Such simulations can help understand how humans may perform movement tasks in different scenarios and could provide useful information in fields like rehabilitation. One of the methodologies used for developing predictive simulations is based on creating models of the neural control architectures that generate the activation of the muscles. In this work we propose the first neural control architecture for cycling behaviors. We show that our architecture can be used to develop predictive simulations of cycling at different speeds and that the associated biomechanical quantitites are consistent with experimental data. Moreover, we show that our control architecture can also replicate walking behaviors with minimal modifications.

## 1. Introduction

Predictive neuromechanical simulations (PNS) have been proposed as a both a way to test motor control theories and as a mean to predict the potential effects of therapy, devices and changes in the biomechanical characteristics of a person [1,2].

PNS are usually obtained from solving an optimal control problem to find the control parameters that allow a biomechanical model to complete a desired task. The biomechanical model represents the human body with an accuracy and physiological constraints that depends on the topic of the research. The solution to the optimal control problem is found through the minimization of the value of a task-specific cost function. Two approaches have been mainly used for both structuring and solving such optimal control problems, direct control and control policy-based control. In the former case the optimal control problems are solved to directly find the states and controls of the model. The most common method to derive the trajectory of the states and controls is Direct Collocation [3]. In control-policy based PNS the optimal control problem is set to find the parameters of a control policy. This control policy consists of a set of physiologically-inspired ruling equations which derive the activations of the actuators of the model. The parameters of the controller are usually found using evolution strategies [4].

Literature presents several examples of predictive simulation studies of cycling. A few seminal studies have used dynamic optimization and genetic algorithms to estimate timing and magnitude of torques or muscular activation that can replicate cycling behaviors [5–7]. Recent studies employ Direct Collocation to investigate different aspects of cycling optimality [8–13], either by tracking experimental data or in fully predictive scenarios. To our knowledge, only few previous works have evaluated the use of control policies for cycling simulations [14,15]. However, these controllers are based on simple biomechanical rules, leaving the full potential of this approach unexplored.

Control-policy based approaches have been widely used to investigate the underlying control of gait. These policies are based on assumptions on neural control of gait and can be classified into reflex-based, CPG-based, or hybrid CPG-reflexes approaches. Each of these approaches can give successful simulations of physiological gait. Reflex-based controllers [16–18] implement feedback reflex networks to drive motion strictly according to sensory input. CPG-based architectures [19] implement central pattern generators [20] as feedforward control that efficiently organizes the model activity. Recently, hybrid controllers, which include both feedback and feedforward components of control, have been proposed to display advantageous properties of both approaches [21,22]. However, a recurrent problem of control policies applied to gait, or other motions, is the impossibility to generalize the controller to other motor tasks. Recent efforts in developing new architectures for PNS have offered breakthroughs in control generalization. For example, a recent work [23] was able to integrate standing up and walking behaviors in a single simulation by attaching sequentially two dedicated controllers. We recently used a modular model to replicate a transition from standing to walking [22].

Here we propose a hybrid neural controller which architecture combines a reflex network and CPGs (**Fig. 1**). The feedback loops represented by the reflexes are physiologically inspired while the CPGs provide a feedforward control which is plausible and it is based on previous successful results [21,24]. This controller aims at representing the first instance of a physiologically-inspired neural controller for cycling. Exploring hypothesis about the underlying neural control of cycling is crucial as it is an exercise often used in the rehabilitation setting. The feedforward component of the controller is inspired by the Unit Burst Generator model proposed by Grillner [25]. In this scheme, CPGs are independent units of bursting activity intertwined in a network. The output of the units depends on their intrinsic properties but also on the configuration of the network. The proposal of this control policy, once implemented, has resulted in the replication of cycling behaviours at different pedaling speeds, presenting metrics that are generally consistent with human data. Additionally, the controller can still present good results when the reflex network is switched off, possibly providing insights on the role of reflexes during cycling. Moreover, we show the potential of this architecture to generalize, with minimal task-specific modifications, to walking behaviors.

**Fig. 1.**
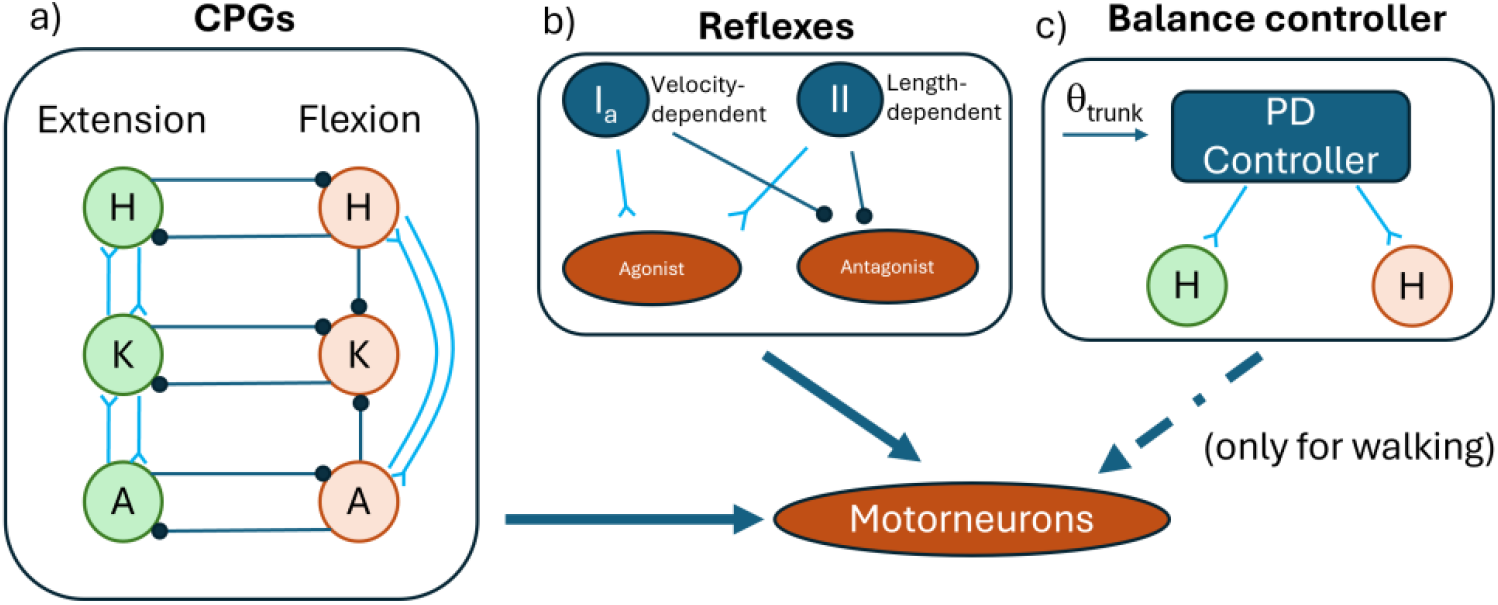
Schematic representation of the controller. The input to the motorneurons is given by the sum of the contributions of the CPGs (a), Reflexes (b) and, for the walking simulations only, the balance controller (c). The CPGs are designed based on the UBG model proposed by Grillner [20]. The reflex architecture is constituted by Ia and II receptors that excite the same muscles and inhibit their antagonist based on velocity and length feedback. The balance controller is a PD controller that regulates the activity of the hip flexors and extensors based on the trunk angle [16].

After showing that this controller can achieve realistic cycling and walking behaviors, we conclude that the combination of reflex and UBG networks offer new and interesting possibilities to develop general control architectures for repetitive lower limbs movements. Such architectures have the potential to be used for personalizing patient-specific control characteristics and evaluate clinically informative predictive simulations of movement for diagnosis and treatment.

## 2. Results

The proposed hybrid controller was able to achieve both cycling and walking behaviors, with biomechanics and muscular activations consistent with those recorded experimentally in both tasks. The joint angles obtained in the simulations mostly agree with the experimental results (**Fig. 2**). We observed a slight over-flexion of the hip at the beginning of the cycle and a slight over-extension of the knee at the dead bottom part of the cycle (50% of the cycle). The ankle angle was consistent with the average values observed experimentally, although an excessive plantarflexion was observed in the second half of the cycle (upstroke) for the 60 and 75 RPMs simulations. These differences at the knee and ankle may be explained, other than from possible limitations of the controller, by the fact that the simulations are in 2D. In fact, in the experimental data the oscillation on the transverse plane may contribute to the joint angle patterns in the sagittal plane.

**Fig 2.**
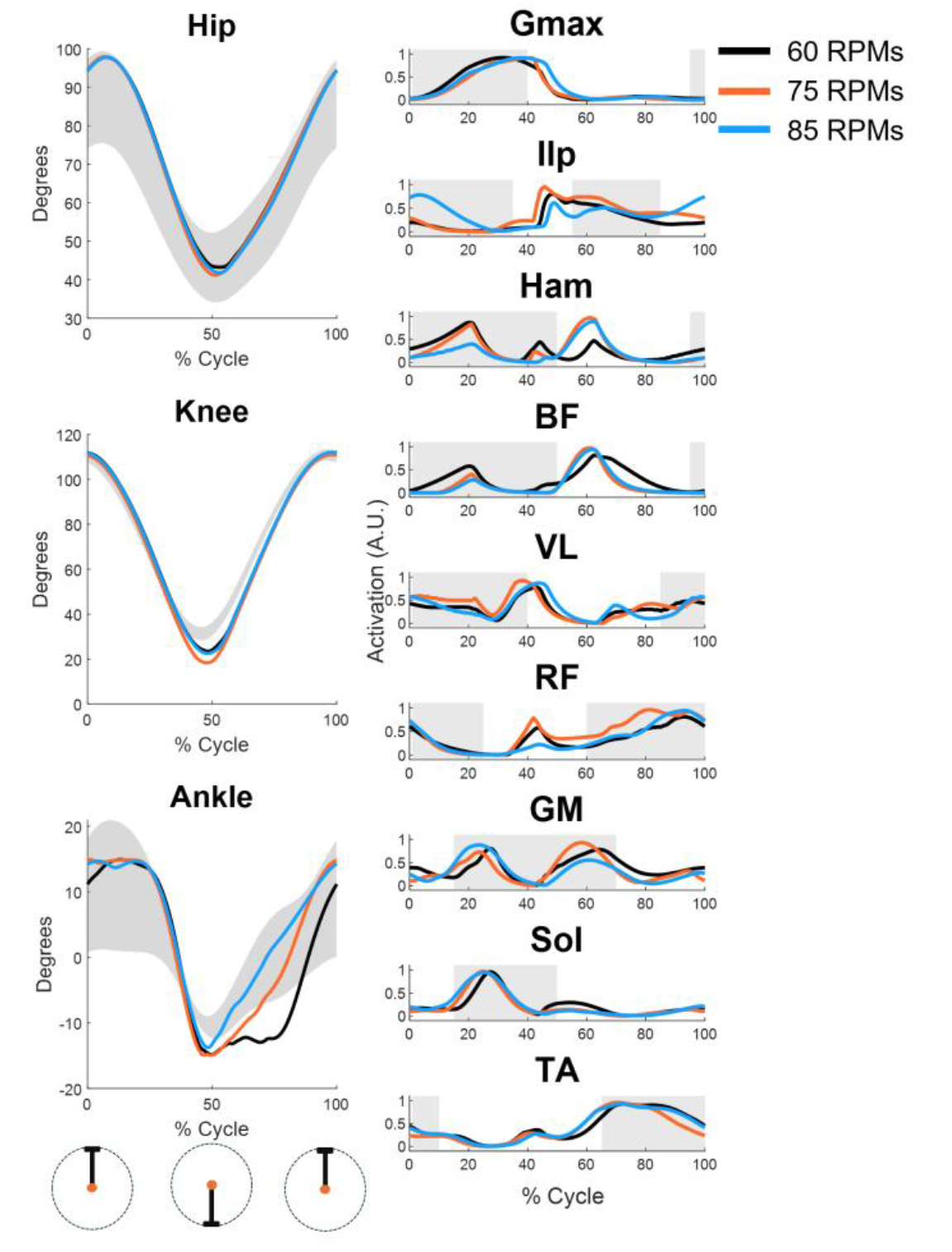
Joint angles (left) and muscular activations (right) obtained from the cycling simulations at 60 (black), 75 (orange) and 85 (blue) RPMs. The

The timings of the muscular activations were compared with on-off dynamics observed experimentally. We observed a general accordance between the simulated and experimental timing of activation of the muscles. HAM and BF displayed patterns which are not entirely physiological. Both presented additional activations during the second half of the cycle. Although consistent with a pulling motion, these activations are not normally observed during standard, non-efficient cycling [26]. We also observed peaks in the activity of the RF, VL and Ilp close to the dead bottom center of the cycle, which are not usually observed during standard cycling. Only minimal differences in muscular activations across the different speeds were observed, with the simulations at 75 and 85 RPMs characterized by a more prominent activity of the HAM during the second half of the cycle.

When comparing the results on cycling between the controller with and without reflexes (**Fig.3**), we observed no substantial differences in the hip and knee angles. Interestingly, the ankle showed a smaller degree of plantarflexion during the second half of the cycle. The muscular activations of the controller without reflexes showed simpler dynamics, characterized by a unique bell-shaped activation. As exception, the VL, SOL and GM presented two peaks. In this controller, the unexpected peaks close to dead bottom observed in the RF and Ilp disappeared, indicating that those spurious peaks depend on the activity of the reflexes.

**Fig 3.**
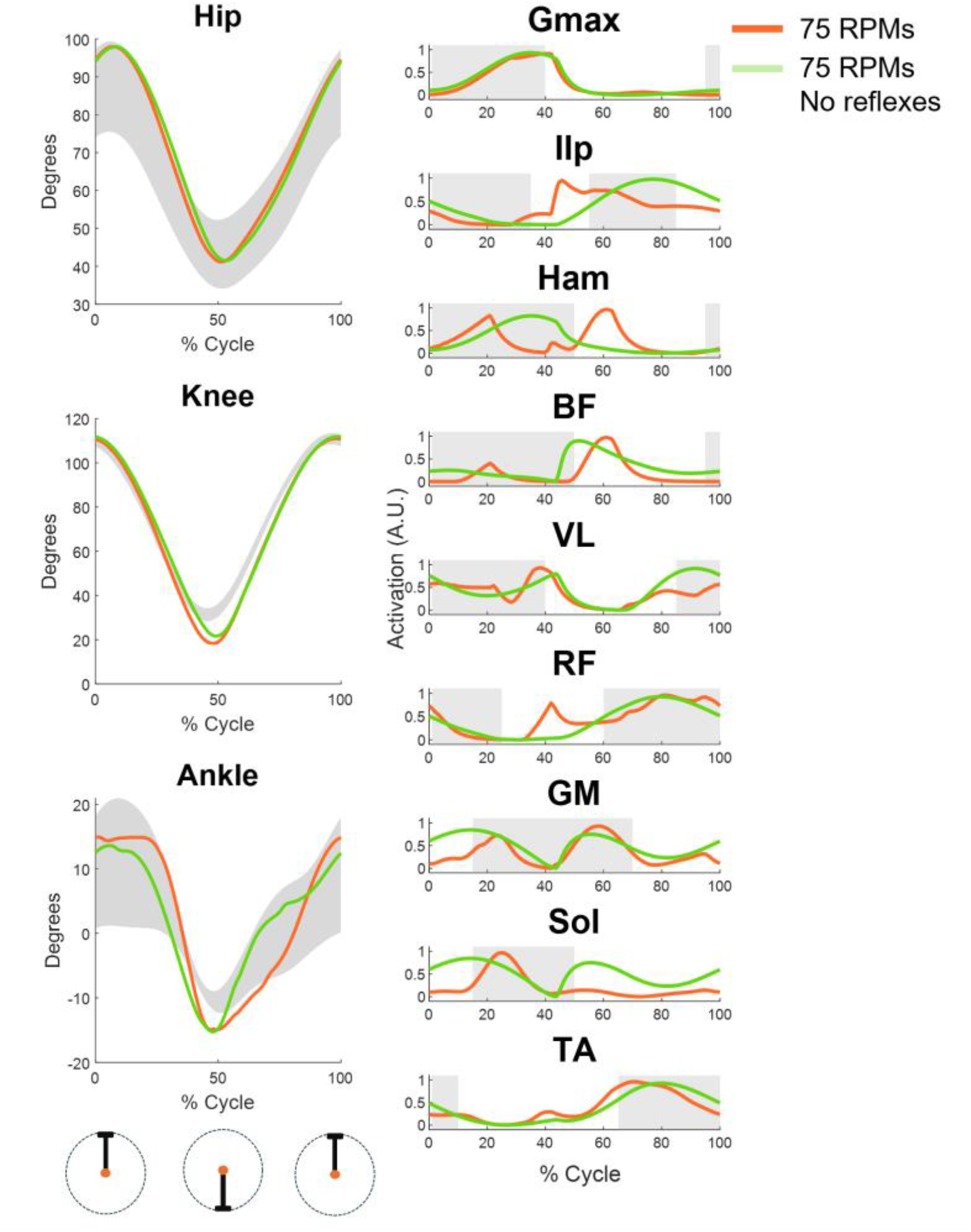
Joint angles (left) and muscular activations (right) obtained from the cycling simulations at 75 RPMs with (orange) and without (green) the reflexes

The most striking differences when the reflexes are absent come from the observation of the Ham and BF muscles. These muscles, in fact, share the same activation dynamics in the full controller, characterized by a physiological peak during the first half of the cycle and a non-strictly physiological peak during the second half. However, when the reflexes are absent, the first peak is only present in the HAM muscle, while the second is only present in the BF (although anticipated). It appears that, for the HAM muscle, the peak during the first half of the cycle is associated with the activity of the hip extensor CPG, as the Gmax is also presenting the same activation pattern. On the other hand, the activity of the BF is only associated with the knee flexion CPG. This is plausible as its peak is consistent with the one observed in GM, which is also a knee flexor, but it is not with HAM, which is also associated to this function. The activation observed in the second half of the cycle in the GM is also consistent with the ankle plantarflexion CPG, given the similarity with the SOL activation, which do not contribute to knee flexion. This latter observation suggests that the ankle plantarflexion and knee flexion CPGs are synchronized during the second part of the cycle.

When optimized for gait the controller achieved a gait speed of 1.46 m/s. The simulations showed physiologically consistent biomechanics and muscular activations, with some small differences (**Fig. 4**). We observed an increment in the hip flexion during late swing, coupled with a slight increased knee flexion and ankle plantarflexion. The ankle pattern lacks a transition from plantarflexion to dorsiflexion during late swing, indicative of a lack of foot preparation for landing. This likely translated in the increased ankle plantarflexion during heel-strike. We also observed an anticipated ankle dorsiflexion during late stance.

**Fig 4.**
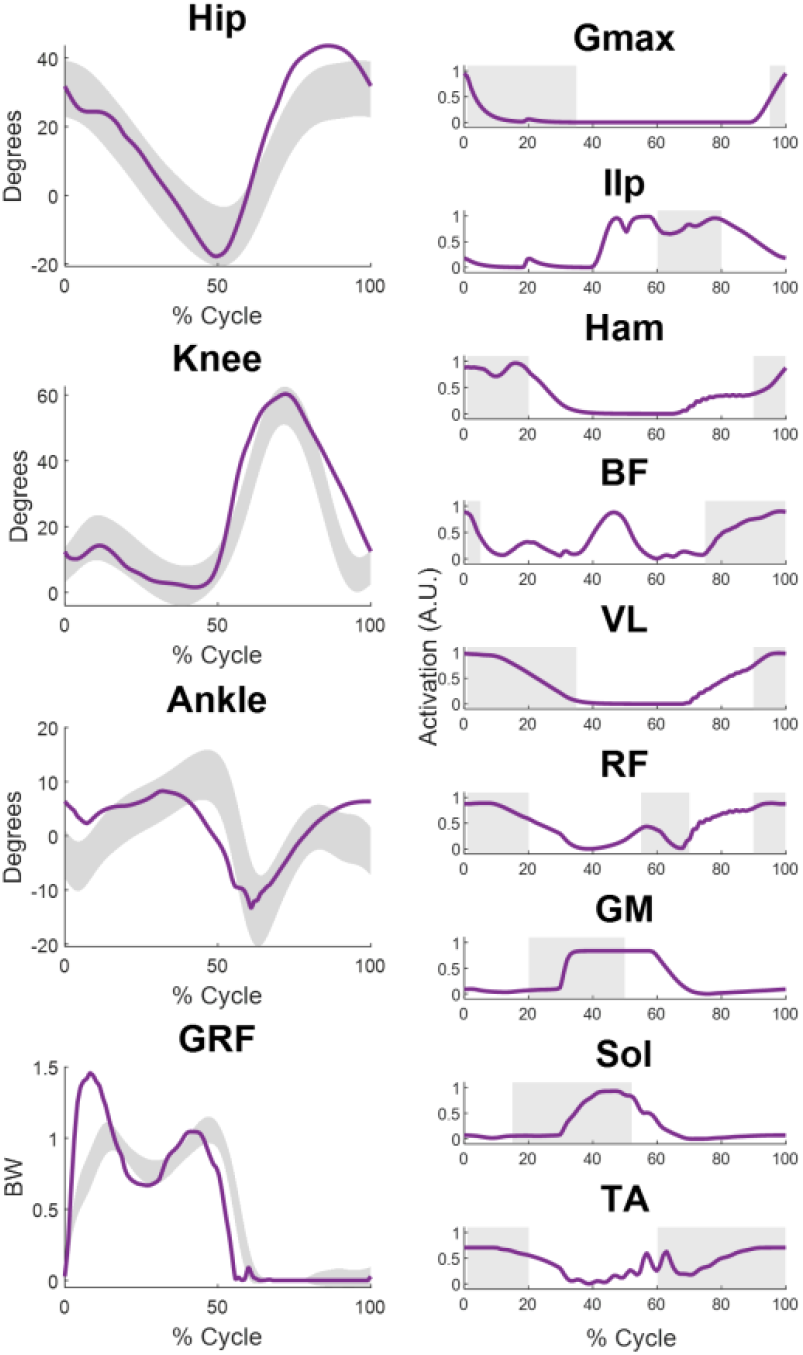
Joint angles (left) and muscular activations (right) obtained from the walking simulations.

The vertical ground reaction forces were consistent with physiological data but characterized by a high peak during load acceptance. This also could be an indicative of a lack of foot preparation during landing. The patterns of muscular activations are generally consistent with the timing observed in experimental studies, with some differences. Such is the case of Ilp, which activation is prolonged, the activation of BF, which presents an unexpected peak during late stance, or delayed activations of the ankle extensors.

We extracted the muscle synergies from both the cycling and walking simulations (**Fig 5**). We used an objective criterion for the selection of the number of synergies [27] which found five synergies for both cycling and walking. Some of the synergies showed a high similarity between the two scenarios. Both tasks presented an almost identical synergy. For example, S5 presented a similarity, as indicated by the cosine product, of 0.93 characterized by the activity of the Ilp, TA and RF. The same three muscles mostly characterized another synergy, S1, that was remarkably similar between the two tasks (similarity of 0.83). In opposition, this synergy is presenting an additional activation of the Gmax for cycling and the Ham for walking. S3 presented a similarity of 0.7 between tasks, arising from the co-activation of the Gmax and VL in both tasks. However, the same synergy also couples the activation of the knee flexors and the TA and RF in walking. S4 represents, for both tasks with a similarity of 0.73, the activity of the ankle plantarflexors, although in cycling we also observe a co-activation of the Gmax. Finally, S2 is the most dissimilar module between the two tasks (similarity of 0.51), sharing only the activity of the BF. Overall, the biggest differences across the modules appear to be related mostly to the activity of the hip flexors and extensors. These muscles are the only which do not share the same control architecture between the two tasks, given the presence of the balance controller for walking.

**Fig 5.**
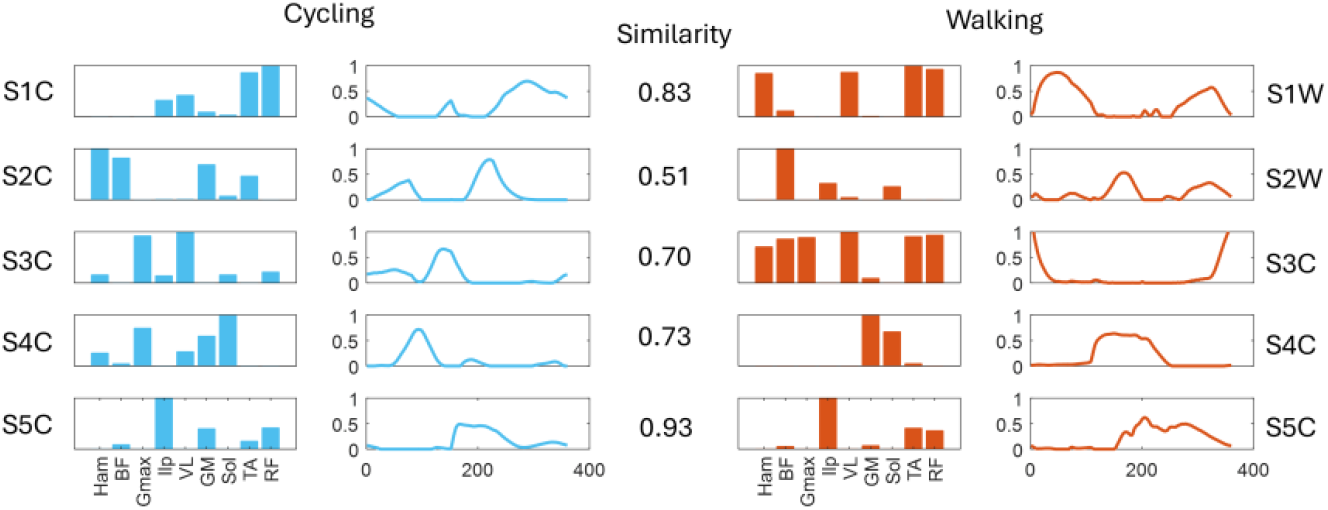
Muscle synergies extracted for cycling (left, blue) and walking (right, red), matched by similarity (calculated as the cosine product between the modules)

## 3. Discussion

In this study we proposed a physiologically plausible hybrid CPG-reflexes neural controller architecture. We showed that, after task-dedicated optimization of the control parameters, the controller can achieve realistic biomechanics and muscular activations during both cycling and walking simulations. The control architecture we propose is one of few in the field of predictive simulations of movement that can generalize across different tasks (Muñoz et al., 2023).

Our work is the first to propose a physiologically plausible neural controller for control-policy based cycling simulations. Recently, Clancy and colleagues developed a DC-based, data-informed simulation of cycling using MOCO [8,28]. In their work, the authors used experimental kinematic and kinetic data to estimate joint, muscle forces and muscular activation in a sample of healthy individuals. Although both works showcase the potential of the PNS for cycling, they also highlight, in our opinion, some conceptual limitations of predictive modelling.

In both works, the simulations result in muscular activation for the hamstrings restricted to upstroke. Although optimal for the task, this dynamic is almost never observed physiologically during standard cycling. The hamstring, in fact, are normally activated in the last phase of downstroke during cycling, while only trained cyclists, by employing a pulling strategy, display hamstrings activation in the first phases of the upstroke [26]. The hamstrings, as knee flexors, when activated during down-stroke, de-facto resist the motion, while their activation during upstroke would provide positive torque towards the cycling motion. This “paradoxical” [29] activation of the hamstrings during cycling is well known and most likely mutuated from their role in controlling final foot position during walking. Predictive simulations of cycling, on the other hand, appear to discover the task-optimal activations. However, as recently pointed out in a review on predictive modeling [2], human neuromuscular control, although well explained by the solution to optimal control problems, is inherently suboptimal. Simulating cycling using optimal control further highlights this suboptimality.

Locomotion and cycling share similar control characteristics. Locomotion, however, is a key human behavior that has been adapted through evolution [30] and whose basic control patterns are observable from early age [31]. Cycling, on the other hand, is a relatively recent behavior. Given its similarities with gait, cycling most likely employs similar control strategies. These strategies, although mostly optimal for gait, may be suboptimal in cycling. An example of this could be the activity of the hamstrings during downstroke. While optimal in the equivalent phase in walking, this activity is suboptimal in cycling. Even though optimal control techniques have shown remarkable potential when simulating human behaviour, task-specific suboptimality needs to be accounted for. Suboptimality can be represented by either adding constrains at the control level or by designing specific cost functions.

We show that the “paradoxical” knee flexor activations during cycling are explained by the reflexive activity included in the cycling controller. The role of reflexes in a hybrid controller of gait has been comprehensively described by Di Russo and colleagues [21,32]. In that work, the authors developed a detailed reflex architecture which uses interneurons regulating the activation of disynaptic reflexes. They also include, differently than this work, reflex pathways controlling excessive force production and muscular activations. Here, given the high number of parameters to be optimized in the CPG controller, we decided to employ a simpler reflex architecture. Such architecture only consisted of monosynaptic equivalents of Ia and II reflex pathways, which regulate activation of the muscles depending on length and velocity feedback. Moreover, we speculated that since we planned to simulate behaviors at submaximal force production condition, where reflexes controlling excessive force production and muscular activations would ideally not be activated, those components may not be needed. The results on cycling without the reflexive components of the controller show that the cycling behavior can be achieved using only the CPGs. Similar results were obtained in the past also for gait using CPG-based control architectures [19,33]. This result opens the question on whether a simple reflex architecture like the one we proposed is necessary during cycling simulation.

The literature on reflexes in gait models, either as sole control components or as sub-components of a hybrid controller, suggest the necessity of including fast feedback pathways in gait simulations [16,21,34–36]. Similarly, the reflex pathways that we included and excluded in our controller are physiologically known to be active during walking [37–40]. Based on the similarity in control between cycling and gait and on the previous argument on how, evolutively, cycling may have control mutuated from walking, it is fair to assume then that reflex pathways are active also during cycling. This has been confirmed also in experimental studies [41,42]. Thus, the potential limitation of the reflex architecture we here propose could be its simplicity rather than its presence.

Our results show that our controller, which was originally designed for cycling, generalizes also to simulating walking behaviors, after adding a control component accounting for balance [16]. The main limitation of the controller during gait appears to depend on the lack of foot preparation for landing during the swing phase. This is a problem that is common to other hybrid controllers (Muñoz et al., 2023; Russo et al., 2023) that do not include state machine-like control during late swing. In our simulations the ankle remains dorsiflexed right before heel strike, rather than returning to a neutral position. It is possible that this behavior may be caused by the direct excitatory link between the hip and the ankle flexion CPGs that we implemented here based on the previously tested UBG-model (Grillner, 20111985; Grillner & Kozlov, 2021). We speculate that including a dynamic balance element in the cost function may lead to better foot-adjustment during swing in our architecture. Dynamic balance have shown to be a major driver of short-term adjustments in gait [43,44]. Proper foot adjustment, in fact, minimizes impact forces and sudden fluctuations in the velocity and acceleration of the center of mass and in the margin of stability.

Cycling and walking have been shown to share similarities in their associated muscle synergies [26,45]. Here we factorized the muscular activity of both tasks into 5 synergies using an objective algorithm [27]. We show that 4 out of 5 of the synergy modules extracted during the two tasks share remarkable similarities. The synergies themselves, compare mostly favourably with experimental synergies observed in literature. For example, S1C, including RF and TA, and S2C, which includes the knee flexors and the Gmax, have been observed in several other experimental works on cycling [26,45,46]. S1W is associated with the activity of the Vasti and has also been observed in experimental studies [45,47,48] while S2W does not present evident similarity with experimentally observed modules. S3C is characterized by the Gmax and VL. However, while experimental studies have shown synergies in cycling where these two muscles contribute together, there is no observation of synergies mostly constituted by these two muscles. The S3W synergy shows activity of the TA, RF and knee flexors. Being active during late swing/early stance, this synergy contributes to response to impact and landing preparation, that, has we already argued, is deficient in our results. In experimental works, these functions are usually disentangled into two synergies [49]. S4C and S4W are the dorsiflexion synergy which is often observed experimentally in both cycling and walking [26,47,50]. Finally, S5C and S5W, which are the most similar between the two tasks, include the hip flexor and the RF. This synergy is commonly observed in walking and has been observed also in cycling, when the hip flexors are included in the recordings [46].

Most of the differences in the synergy modules between the two simulated tasks is accounted by differences in the activation of the Ham (S1-3C and S1-3W), BF (S3C and S3W) and Gmax (S4C and S4W). This could be expected as these are the only muscles with differences in the control architecture, given the presence of the balance controller for gait. The balance controller is a PD controller active during stance that regulates the activity of the hip muscles based on feedback on the trunk position. The presence of feedback components regulating the angle of the trunk with respect to the pelvis during walking is plausible, although their exact architecture is not known. Thus, we cannot exclude that a physiological balance controller would be constituted by feedback-based circuits mapping directly at the level of the CPGs rather than on the single muscles. Synergies have often been indicated as the building blocks of movement and believed to reflect networks of spinal interneurons coordinating the activity of muscles in group [51]. Our results show that physiologically plausible synergies arise from the coordinated activity of flexion and extension CPGs and reflexes. In the upper limbs, synergies can be easier to analyze as coordination encoded at the spinal level which do not depend on reflexes and cyclic activation. In opposition, our results show that lower limb synergies, obtained from factorizing EMG activity, may mask different control pathways contributing to the generation of the muscular activations.

Predictive simulations have long been proposed as a mean to test the effect of therapies or changes in characteristics of impairment [1]. The few works that have tested the used of PNS of impairment were mostly based on DC-based simulative architectures [52–54] Recently, control-policy based methods have been used on biomechanical models simulating impairments [55]. While DC-based simulations show remarkable potential in the simulation of steady-state behaviors, the incorporation of feedback is still being investigated [56]. DC-based simulations offer little possibility of personalization in the control strategy, thus limiting the possibility of making patient-specific models. Control personalization has, so far, only been modeled as patient-specific muscle synergies, or through the modification of the neuromechanical characteristics of the biomechanical and muscle model to account for deficits such as spasticity or weakness [52,57]. Here we show that synergies can arise from the activity of CPGs and reflexes, and do not appear to differentiate between the different control components. Conditions such as stroke can affect both feedback and feedforward components of control, differently. Stroke survivors often exhibit hyperreflexia, that can be modeled in the feedback component of a control-policy based architecture (as shown in a recent work on Cerebral Palsy [58]) but also in the simplified control of the feedforward component. Given this rationale, control-policy based predictive simulations offer, at least theoretically, greater possibility of control personalization and could be effectively used to model the effect that different intervention may have on the different control components, separately. The drawback, however, of policy-based simulation is the impossibility to validate the control underlying the simulation as accurate [59]. For this reason, developing control policy that are fully based on experimentally validated principles, although not a validation, represent the safer approach. Our work presents some limitations. Both the biomechanical model and the controller are simple when compared with physiology. The reflex architecture is simple with respect to current models [16,21]. Our CPG implementation does not account for phase resetting. Moreover, we completely omitted possible inhibitory and excitatory connections, both in the reflexes and the CPGs, between the two legs. Additionally, our simulations are constrained to a 2D scenario. While this simplicity limits the comparison of our results with experimental ones, it is dictated by the necessity of testing the feasibility of the proposed approach. Although the controller presents a reductive reflex architecture, and it was applied to a simple biomechanical model already tested [16,21], it needed a multi-staged optimization of more than 100 parameters. Thus, optimization time, difficulties in convergence and risk of genetic drift [60] increased. Another limitation of our work, shared with similar works in literature, is the empirical nature of the cost functions that we employed. The weights associated with the cost function elements were selected empirically based on preliminary results. A more substantial analysis of the elements of the different cost functions and their associated weight, also considering different weighting techniques [61], would likely improve our result.

## 4. Methods

### 4.1 The Biomechanical model

The cycling and gait simulations were based on the same 2-D biomechanical model, applied to the two different scenarios (**Fig. 6**). The model was derived from the one developed by Delp and colleagues [62] which was also used in similar works [16,21]. Seven segments represent the human body in this biomechanical model. Feet, shanks, thighs and pelvis are present in the cycling scenario, while feet, shanks, thighs and HAT (head-arms-torso) are present in walking. The model presents 9 DoFs for the gait environment, consisting of pelvis tilt and anteroposterior-longitudinal translations, and hip, knee and ankle frontal rotation. In the cycling environment, pelvis tilt and translations were fixed, and a gear-crank-pedals assembly was attached to the model. This implementation of the model also presents 9 DoFs, consisting of gear and pedals rotation, and hip, knee and ankle frontal rotation. The height of the seat in the cycling model was set to match the criteria for seat height used in Clancy et al. [8]. The model, in both scenarios, present 18 Hill-type muscles (9 per legs): hamstrings (Ham), biceps femoris short head (BF), gluteus maximus (Gmax), iliopsoas (Ilp), vastus medialis (VM), rectus femoris (RF), gastroecnemius medialis (GM), soleus (Sol) and tibialis anterior (TA). The model was implemented in the SCONE software [63] for the Hyfydy simulation engine [64].

**Fig. 6.**
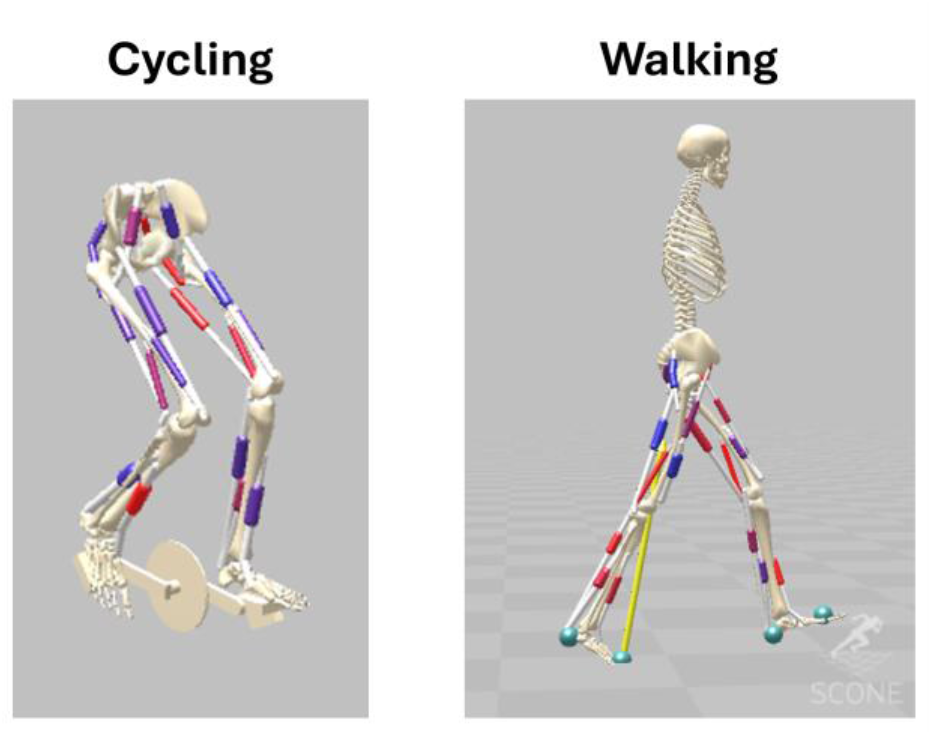
The biomechanical models for cycling used in the simulations

### 4.2 The controller

The controller was designed as a hybrid controller composed by components of feedforward and feedback control. The feedforward component is based on the Unit Burst Generator model (UBG-model) proposed by Grillner [20] (**Fig. 1a**). This model consists of unit CPGs inter-connected in a network. The feedback component of the controller consists of a set of reflexes regulating the muscular activity based on sensory inputs. Particularly, muscle length and velocity feedback reflexes (**Fig. 1b**) are implemented in the controller. For the walking simulation, a third component, a balance controller (**Fig. 1c**), is added. This controller is an additional feedback component which is active during early and late stance. The role of this controller is to control the trunk angle by acting only on the hip muscles. This control has been used in previous models [16,21,35]. For each muscle, the excitation at each instant is the sum of the excitation arising from all CPGs and reflex pathways in which the muscle is involved. Hip muscles receive also an excitatory component from the balance controller during the walking simulations.

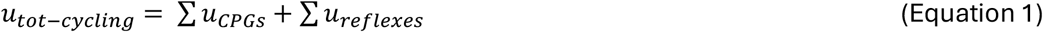

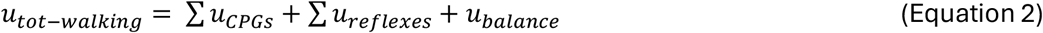

where *u*_*tot-cycling*_ and *u*_*tot-walking*_ are the total output of the controller to generate muscular activity for cycling and walking behaviors, respectively, *u*_*CPGs*_ is the output of the UBG-model, and *u*_*reflexes*_ is the output of the reflex network. The component *u*_*balance*_ is the output of the balance controllers, only employed for walking simulations.

### 4.3 The Central Pattern Generators

The CPG architecture is based on the previously proposed UBG-model [25]This model is constituted by unit CPGs which control a set of motoneurons [65]. The units are distributed in a network which consists of inhibitory and excitatory synapses. This coupling between units, besides their intrinsic properties, determines their output.

In the 2D model we employed, each unit is controlled by a rhythmic activation that acts as a clock function. We identified a total of 12 units, 6 per leg, consisting of flexor and extensor units acting on the hip, knee and ankle. In the overall network the different units act as reciprocal inhibitors and/or excitators for the other units, following a scheme precedently proposed in a simulation study [66], and based on experimental observations [67]. In this scheme (**Fig. 2a**), pairs of flexor and extensor units acting on the same joint provide reciprocal inhibition. Extensors units provide reciprocal excitation to the units of the closest joint (e.g. hip extensors excite knee extensors, which excite both hip and ankle extensors). Flexors units follow a more complex inhibition/excitation pattern, with the hip and ankle units inhibiting the knee unit and exciting each other. The activation of each single unit, as already proposed previously [21], is based on raised cosine function:

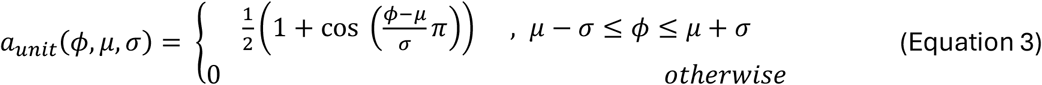

where *ϕ* is the leg-dependent gait phase, *μ* is the position of the peak of the activation in the cycle, and *σ* is the witdth of the activation pattern. The output, a_unit_, is expressed in the form of a discrete bell-shaped waveforms, in opposition to a classical oscillator configuration [68]. Differently by the well-known Aoi’s model of dynamically coupled CPGs [24], where the phase generator for both sides is governed by a differential equation coupling the two legs, here we fixed the coupling between the two legs across the whole simulation. Moreover, while previous implementations [21,24] used the heel-strike and foot-off to reset the phase, we did not add an event-based phase reset. Instead, the phase continuously increases. These choices were made under the assumption that our simulations will replicate steady-state unperturbed cycling and walking, where the coupling of the two legs is always maintained to a delay of half-cycle. Thus, at the beginning of each simulation we set *ϕ*_*left*_ = 0 *and ϕ*_*right*_ = *π*. The phases are then updated at each instant following the equation:

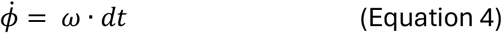

where *ω* is the angular frequency common to all the 12 oscillators and *dt* is the time-step of the simulation. This equation establishes the phase generator, or clock function, of the system. The activation of each single unit was then calculated based on the UBG architecture as follows:

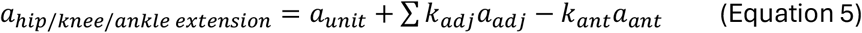

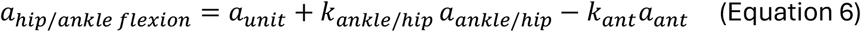

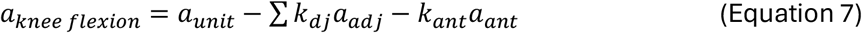

Where the *a* parameters are the outputs of the different unit CPGs and represent the *u*_*CPG*_ values from equation 1. The *k* parameters are weights in the interval [0;1] whose values are set through the optimization process. The suffixes *adj* and *ant* in the *a* and *k* parameters in equations (5-7) mean adjacent and antagonist respectively. For the extension CPGs, each unit has a total output that is the sum of the activation of the unit itself, plus a weighted excitatory contribution coming from the adjacent units and a weighted inhibitory contribution coming from the corresponding antagonist flexion unit. For flexion, hip and ankle receive a weighted excitatory contribution from each other and inhibition from the corresponding extension unit. The flexion knee receives inhibition from all adjacent units and the corresponding extension unit. The contribution of the units to each muscle (*u*_*muscle/CPG*_) is calculated as:

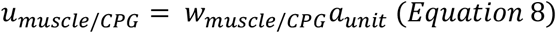

where the *w* parameters are weights regulating the contribution of each muscle associated to a unit. The muscles only receive activity from the units to which they are associated. For example, Gmax only receives the output generated by the hip extension unit, while RF receives output from both the hip flexion and knee extension units. The feedforward component of the controller has a total of 37 parameters to be optimized. Those parameters include the weights *w* and *k*, the values of *μ* and *σ* of each CPG, and the general angular frequency of the clock function *ω*. To minimize the number of parameters to be optimized, we assumed complete symmetry between the two legs and the same values were associated to parameters mapping on the right and left leg.

### 4.4 The reflex architecture

In our control architecture we employed a series of reflex pathways, mutuated from previous works [21,69] but simplified based on task-related considerations. The reflex architecture we employed include:

a. Ia afferents, providing self-excitation and agonist inhibition to the muscles based on a velocity-dependent response to stretch [40].
b. II afferents providing excitation to the same muscle and inhibition to the antagonist muscle(s) based on muscle length [39].

We did not include Ib afferents and Reshaw cells in our architecture [37,38]. Ib afferents provide same and antagonistic inhibition and are triggered by large forces. Renshaw cells provide same and antagonistic inhibition and are triggered by excessive muscular activations. Here we perform simulations on two tasks, cycling and walking, both performed at a sub-maximal exertion level and under an energy minimization constraint. We then assume that, under these task conditions, Ib afferents and Renshaw cells, which are both mechanisms controlling excessive force production, are not expected to contribute significantly. The reflexes were all modeled as monosynaptic reflexes, differently from the comprehensive interneuron-mediated network presented by Di Russo [21]. This choice was made to minimize the number of parameters to be optimized. The excitations relative to the reflexes were calculated following equations 9 and 10 for Ia and II afferents respectively:

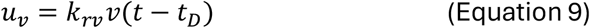

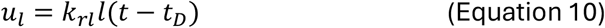

where *k*_*rv*_ and *k*_*rl*_ are the gains of the reflexes, *v* and *l* are muscle velocity and length, and *t*_*D*_ is the muscle-specific time delay, 5, 10 or 20 ms, determined from previous studies on reflex-based controllers [16]. The gains *k* are set during the optimization procedure and are positive for excitatory reflexes and negative for inhibitory ones. Each muscle is characterized by 2 excitatory gains relative to Ia and II, and one or two inhibitory gains to its antagonists. In total, 46 gains (18 excitatory, 28 inhibitory) were optimized during both gait and cycling. Weights were also assumed symmetrical between the two legs to minimize the number of parameters to be optimized.

### 4.5 The Balance controller

In the walking simulations, due to the need for stability, a balance controller was added to the architecture. This component consists of the trunk control policy originally implemented by Ong et al. [16]. The balance controller is implemented based on a state-machine and is only active during early and late stance. The objective of this new component is the regulation of the trunk angle by acting on the Ilp, Gmax and Ham muscles. The controller is a PD controller and its output follows the equation:

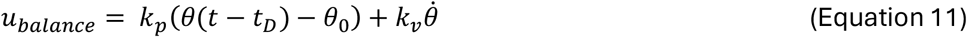

where *k*_*p*_ and *k*_*v*_ are the proportional and derivative gains, *θ* is the trunk angle (to which a time delay equal to 5 ms is applied) and *θ*_0_ is the desired value for the trunk angle. A total of 9 parameters were optimized for this controller.

### 4.6 Optimization procedures

The cycling and walking simulations had 101 and 110 parameters to be optimized, respectively. The parameters are divided in 18 position and velocity offsets to the initial pose, 37 CPG parameters, 46 reflex parameters, and the additional 9 parameters of the balance controller only present for the gait simulations. The optimization was performed using the Covariance Matrix Adaptation Evolutionary Strategy, implemented on the SCONE software [63]. Due to the number of parameters to be optimized we employed a multi-stage optimization procedure where the parameters found in one stage were used as the initial guess for the following stage.

### 4.7 Cycling task optimization

The optimization for cycling was divided in two stages. A first stage found the initial cycling solutions that matched the target speed of pedaling while limiting the range of motion of the ankle. The second stage, employing the solution of the first stage as initial guess, aimed at finding parameters that minimized energy expenditure and the loads at joints, while limiting their range of motion to physiological values. Additionally, solutions with inactive muscles were penalized. The cost function for the stage 1 was then:

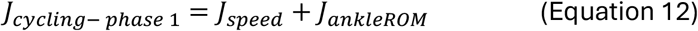

where the element *J*_*speed*_ minimizes the difference between the desired speed (set at 60, 75 or 85 RPMs) and the actual speed achieved by the model during the simulation, while the element *J*_*ankleROM*_ restricts the ankle range of the solution to a physiologically observed range of [-15 deg; 15 deg]. The cost function for the stage 2 was:

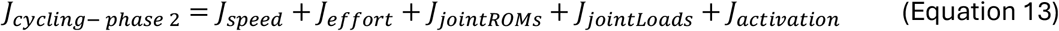

where element *J*_*speed*_ is equivalent to the one used in stage 1; the element *J*_*effort*_ minimizes the sum of the squared muscle activations of all the muscles; element *J*_*jointROMs*_ is the extension of element *J*_*ankleROM*_ of stage 1 to all the joints, restricting the ranges of the hip, knee and ankle joints to [25 deg; 95 deg], [25 deg;120 deg] and [-15 deg; 15 deg] respectively; element *J*_*jointloads*_ minimizes the loads at all the joints, as shown to be beneficial in previous cycling simulations [8]; element J_activation_ penalizes muscular activations below 0.1 in a range [0; 1]. This latter element aims at avoiding solutions where one or more muscles are inactive. A low muscle activation profile is likely to happen during the optimization process as the control architecture employed in this study includes several inhibitory pathways between multiple sub-blocks of control. Each optimization stage was run for a maximum of 10,000 generations.

### 4.8 Walking task optimization

The optimization for walking was divided in three stages, in a scheme similar to the one proposed by Di Russo and colleagues [21]. The stage 1 was not used to achieve a gait behavior. Instead, the objective is initializing the parameters to achieve segment movements and muscular activations similar to those of a physiological gait behavior. The biomechanics and muscular activations to imitate were obtained by running the optimized reflex-based controller developed by Ong and colleagues [16]. The cost function for stage 1 was then:

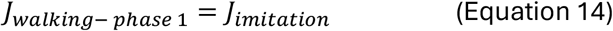

where *J*_*imitiation*_ minimizes the difference of the kinematics and muscular activations obtained from our model respect to the simulation results derived from the Ong controller. The stage 2 of the optimization aimed at obtaining a realistic gait behavior, using the parameters obtained in stage 1. The cost function for stage 2 was:

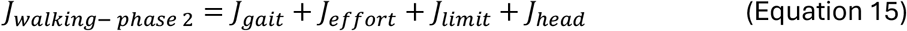

where *J*_*gait*_ penalizes solutions outside the set velocity range (1.5 ± 0.1 m/s) and solutions where the model falls. *J*_*effort*_ penalizes the metabolic cost of walking following the methodology developed by Uchida and colleagues [70]. *J*_*limit*_ minimizes the joint limit torques at the knee and ankle, and *J*_*head*_ minimizes the head acceleration in the vertical and horizontal directions. Finally, the optimization in stage 3 is used to discard solutions with inactive muscles and the cost function is:

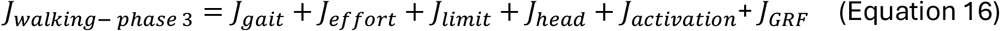

where the first four elements of the cost function are equivalent to stage 2. The *J*_*activation*_ element penalizes muscular activations below 0.1 in a range [0; 1], as in the cycling optimization. Finally, the *J*_*GRF*_ element penalizes peak values of vertical GRF exceeding 1.5 the body weight of the model. All elements of the cost functions in the three stages were associated with a specific weight, manually selected based on previous, unreported, simulations.

### 4.9 Simulations

For the cycling scenario, we performed four sets of simulations. For each set of simulations, we ran ten independent optimizations for stage one, then the result with the best convergence was used as starting condition for stage two. The quality of the convergence was based on the fitness calculated from the cost function. Stage two consisted of five optimization processes. The results presented are, again, the best solution among the five optimizations. The first three sets consisted of simulation scenarios performed using the full controller (CPG and reflexes) at three different target speeds of 60, 75 and 85 RPMs. The speed was modulated by changing the element *J*_*speed*_ in equations 12 and 13. The aim of these simulations was to test the consistency of the results across different speeds. The last set of cycling optimizations was run using only the CPG component of the controller, without the reflex architecture. The aim of this simulation was evaluating the effect of speed and length-dependent reflexes during cycling.

For the walking scenario, we ran a single set of simulations, to evaluate the capacity of the controller to generalize to a different motor task. We ran ten independent optimization processes for stage one and stage two, and five processes for stage three.

### 4.10 Validation

To evaluate the ability of our controller to replicate physiological cycling and gait, we compared biomechanical metrics in both tasks with those recorded in existing accessible datasets. The metrics consisted of the angular positions of the joints, and the muscular patterns of activation. Specifically, for cycling, we used the dataset used in Clancy et al. [8] to have the comparison of the kinematics. The dataset was recorded from 16 healthy individuals cycling on a stationary bike at self-selected speeds ranging from 77 to 92 RPMs. The muscular activations were compared to the activation timing observed in experimental cycling setups similar to our simulation [26,46]. For walking, we used the dataset recorded by Van Criekinge and colleagues [71] for both, kinematics and muscle activations. The dataset was recorded from 138 healthy individuals walking at self-selected speed.

### 4.11 Muscle Synergies Analysis

Previous works in the literature have reported on the similarity between the muscle synergies extracted during walking and cycling [26,45]. We here compared the synergies extracted from our cycling and walking simulations. Our objective is to assess whether the muscular activations obtained by applying the same control architecture to two different tasks result in similar synergy modules. We extracted synergies from the muscular activations derived from the simulations using the standard non-negative matrix factorization algorithm [72]. A previously validated objective methodology was used to derive the optimal number of synergies for each task [27]. The similarity between the synergy modules was calculated using the cosine product.

## Data availability statement

There are no primary data in the paper; All code will be uploaded on Zenodo

## Acknowledgments

The authors like to acknowledge Thomas Geijtenbeek, Magdalena Zych and Ziqi Peng for their help in the development of the biomechanical model used in the simulations and of the code used for running the simulations. This project was supported by Science Foundation Ireland (Grant No. 21/FFP-P/10228).

## References

[1] De Groote F, Falisse A. Perspective on musculoskeletal modelling and predictive simulations of human movement to assess the neuromechanics of gait. Proceedings of the Royal Society B: Biological Sciences 2021;288. 10.1098/rspb.2020.2432.

[2] Bersani A, Davico G, Viceconti M. Modeling Human Suboptimal Control: A Review. J Appl Biomech 2023;39. 10.1123/jab.2023-0015.

[3] De Groote F, Kinney AL, Rao A V., Fregly BJ. Evaluation of Direct Collocation Optimal Control Problem Formulations for Solving the Muscle Redundancy Problem. Ann Biomed Eng 2016;44:2922–36. 10.1007/s10439-016-1591-9.

[4] Dorn TW, Wang JM, Hicks JL, Delp SL. Predictive simulation generates human adaptations during loaded and inclined walking. PLoS One 2015;10:1–16. 10.1371/journal.pone.0121407.

[5] Kautz SA, Hull ML. Dynamic optimization analysis for equipment setup problems in endurance cycling. J Biomech 1995;28:1391–401. 10.1016/0021-9290(95)00007-5.

[6] Van Soest AJK, Casius LJR. Which factors determine the optimal pedaling rate in sprint cycling? Med Sci Sports Exerc 2000;32:1927–34. 10.1097/00005768-200011000-00017.

[7] Bobbert MF, Casius LJR, Van Soest AJ. The Relationship between Pedal Force and Crank Angular Velocity in Sprint Cycling. Med Sci Sports Exerc 2016;48:869–78. 10.1249/MSS.0000000000000845.

[8] Clancy CE, Gatti AA, Ong CF, Maly MR, Delp SL. Muscle-driven simulations and experimental data of cycling. Scientific Reports 2023 13:1 2023;13:1–10. 10.1038/s41598-023-47945-5.

[9] Kistemaker DA, Terwiel TM, Reuvers EDHM, Bobbert MF. Limiting radial pedal forces greatly reduces maximal power output and efficiency in sprint cycling: an optimal control study. J Appl Physiol (1985) 2023;134:980–91. 10.1152/JAPPLPHYSIOL.00733.2021/ASSET/IMAGES/LARGE/JAPPLPHYSIOL.00733.2021_F008.JPEG.

[10] Park S, Caldwell GE, Umberger BR. A direct collocation framework for optimal control simulation of pedaling using OpenSim. PLoS One 2022;17:e0264346. 10.1371/JOURNAL.PONE.0264346.

[11] Jansen C, McPhee J. Predictive Dynamic Simulation of Seated Start-Up Cycling Using Olympic Cyclist and Bicycle Models. Proceedings 2018, Vol 2, Page 220 2018;2:220. 10.3390/PROCEEDINGS2060220.

[12] Jansen C, Mcphee J. Predictive dynamic simulation of Olympic track cycling standing start using direct collocation optimal control. Multibody Syst Dyn 2020;49:53–70. 10.1007/s11044-020-09723-3.

[13] Farahani SD, Bertucci W, Andersen MS, Zee M de, Rasmussen J. Prediction of crank torque and pedal angle profiles during pedaling movements by biomechanical optimization. Structural and Multidisciplinary Optimization 2015;51:251–66. 10.1007/S00158-014-1135-6/FIGURES/9.

[14] Raasch CC, Zajac FE. Locomotor strategy for pedaling: Muscle groups and biomechanical functions. J Neurophysiol 1999;82:515–25. 10.1152/JN.1999.82.2.515/ASSET/IMAGES/LARGE/9K0790328007.JPEG.

[15] Chen G, Kautz SA, Zajac FE. Simulation analysis of muscle activity changes with altered body orientations during pedaling. J Biomech 2001;34:749–56. 10.1016/S0021-9290(01)00014-8.

[16] Ong CF, Geijtenbeek T, Hicks JL, Delp SL. Predicting gait adaptations due to ankle plantarflexor muscle weakness and contracture using physics-based musculoskeletal simulations. PLoS Comput Biol 2019. 10.1371/journal.pcbi.1006993.

[17] Song S, Geyer H. Evaluation of a neuromechanical walking control model using disturbance experiments. Front Comput Neurosci 2017;11. 10.3389/fncom.2017.00015.

[18] Geyer H, Herr H. A Muscle-reflex model that encodes principles of legged mechanics produces human walking dynamics and muscle activities. IEEE Transactions on Neural Systems and Rehabilitation Engineering 2010. 10.1109/TNSRE.2010.2047592.

[19] Taga G. A model of the neuro-musculo-skeletal system for human locomotion - II. Real-time adaptability under various constraints. Biol Cybern 1995;73. 10.1007/BF00204049.

[20] Grillner S. Neurobiological Bases of Rhythmic Motor Acts in Vertebrates. Science (1979) 1985;228:143–9. 10.1126/SCIENCE.3975635.

[21] Di Russo A, Stanev D, Sabnis A, Danner SM, Ausborn J, Armand S, et al. Investigating the roles of reflexes and central pattern generators in the control and modulation of human locomotion using a physiologically plausible neuromechanical model. J Neural Eng 2023:2023.01.25.525432. 10.1101/2023.01.25.525432.

[22] Munoz D, Holland D, Severini G. A novel modular architecture for a neural controller architecture for predictive simulations of stand-to-walk motions. BioRxiv 2023:2023.12.04.569887. 10.1101/2023.12.04.569887.

[23] van der Kruk E, Geijtenbeek T. A planar neuromuscular controller to simulate compensation strategies in the sit-to-walk movement. PLoS One 2024;19:e0305328. 10.1371/JOURNAL.PONE.0305328.

[24] Aoi S, Ogihara N, Funato T, Sugimoto Y, Tsuchiya K. Evaluating functional roles of phase resetting in generation of adaptive human bipedal walking with a physiologically based model of the spinal pattern generator. Biol Cybern 2010;102:373–87.

[25] Grillner S, Kozlov A. The CPGs for Limbed Locomotion–Facts and Fiction. International Journal of Molecular Sciences 2021, Vol 22, Page 5882 2021;22:5882. 10.3390/IJMS22115882.

[26] De Marchis C, Schmid M, Bibbo D, Castronovo AM, D’Alessio T, Conforto S. Feedback of mechanical effectiveness induces adaptations in motor modules during cycling. Front Comput Neurosci 2013;7:42906. 10.3389/FNCOM.2013.00035/BIBTEX.

[27] Ranaldi S, De Marchis C, Severini G, Conforto S. An Objective, Information-Based Approach for Selecting the Number of Muscle Synergies to be Extracted via Non-Negative Matrix Factorization. IEEE Transactions on Neural Systems and Rehabilitation Engineering 2021;29. 10.1109/TNSRE.2021.3134763.

[28] Dembia CL, Bianco NA, Falisse A, Hicks JL, Delp SL. OpenSim Moco: Musculoskeletal optimal control. PLoS Comput Biol 2020;16:e1008493. 10.1371/JOURNAL.PCBI.1008493.

[29] Andrews JG. The functional roles of the hamstrings and quadriceps during cycling: Lombard’s Paradox revisited. J Biomech 1987;20:565–75. 10.1016/0021-9290(87)90278-8.

[30] Duysens J. Human gait as a step in evolution. Brain 2002;125:2589–90. 10.1093/BRAIN/AWF261.

[31] Dominici N, Ivanenko YP, Cappellini G, D’Avella A, Mond V, Cicchese M, et al. Locomotor primitives in newborn babies and their development. Science (1979) 2011;334:997–9. 10.1126/science.1210617.

[32] Di Russo A, Koelewijn AD, Dzeladini F, Ijspeert AJ. Neuromechanical simulation of human locomotion: descending modulation of spinal reflex parameters during speed changes. 9th International Symposium on Adaptive Motion of Animals and Machines (AMAM 2019), 2019.

[33] Aoi S, Funato T. Neuromusculoskeletal models based on the muscle synergy hypothesis for the investigation of adaptive motor control in locomotion via sensory-motor coordination. Neurosci Res 2016;104:88–95. 10.1016/j.neures.2015.11.005.

[34] Geyer H, Herr H. A Muscle-reflex model that encodes principles of legged mechanics produces human walking dynamics and muscle activities. IEEE Transactions on Neural Systems and Rehabilitation Engineering 2010;18:263–73. 10.1109/TNSRE.2010.2047592.

[35] Song S, Geyer H. Evaluation of a neuromechanical walking control model using disturbance experiments. Front Comput Neurosci 2017;11:250076. 10.3389/FNCOM.2017.00015/BIBTEX.

[36] Ramadan R, Geyer H, Jeka J, Schöner G, Reimann H. A neuromuscular model of human locomotion combines spinal reflex circuits with voluntary movements. Scientific Reports 2022 12:1 2022;12:1–23. 10.1038/s41598-022-11102-1.

[37] Neurosciences CC-T in, 2001 undefined. Force-feedback during human walking. CellComC CapadayTRENDS in Neurosciences, 2001•cellCom n.d.

[38] Veale JL, Rees S. Renshaw cell activity in man. J Neurol Neurosurg Psychiatry 1973;36:674–83. 10.1136/JNNP.36.4.674.

[39] Lundberg A, Malmgren K, Schomburg ED. Reflex pathways from group II muscle afferents - 1. Distribution and linkage of reflex actions to α-motoneurones. Exp Brain Res 1987;65:271–81. 10.1007/BF00236299/METRICS.

[40] Chen HH, Hippenmeyer S, Arber S, Frank E. Development of the monosynaptic stretch reflex circuit. Curr Opin Neurobiol 2003;13:96–102. 10.1016/S0959-4388(03)00006-0.

[41] Grey MJ, Pierce CW, Milner TE, Sinkjaer T. Soleus Stretch Reflex during Cycling. Motor Control 2001;5:36–49. 10.1123/MCJ.5.1.36.

[42] Brown DA, Kukulka C. Human flexor reflex modulation during cycling. Article in Journal of Neurophysiology 1993. 10.1152/jn.1993.69.4.1212.

[43] Severini G, Zych M. Locomotor adaptations: paradigms, principles and perspectives. Progress in Biomedical Engineering 2022;4. 10.1088/2516-1091/ac91b6.

[44] Cajigas I, Koenig A, Severini G, Smith M, Bonato P. Robot-induced perturbations of human walking reveal a selective generation of motor adaptation. Sci Robot 2017;2:eaam7749. 10.1126/scirobotics.aam7749.

[45] Barroso F, Torricelli D, Moreno JC, Taylor J, Gomez-Soriano J, Esteban EB, et al. Similarity of muscle synergies in human walking and cycling: Preliminary results. Proceedings of the Annual International Conference of the IEEE Engineering in Medicine and Biology Society, EMBS 2013:6933–6. 10.1109/EMBC.2013.6611152.

[46] Zych M, Rankin I, Holland D, Severini G. Temporal and spatial asymmetries during stationary cycling cause different feedforward and feedback modifications in the muscular control of the lower limbs. J Neurophysiol 2019;121:163–76. 10.1152/jn.00482.2018.

[47] Zych M, Cannariato A, Bonato P, Severini G. Forward and backward walking share the same motor modules and locomotor adaptation strategies. Heliyon 2021;7. 10.1016/j.heliyon.2021.e07864.

[48] Severini G, Koenig A, Adans-Dester C, Cajigas I, Cheung VCK, Bonato P. Robot-Driven Locomotor Perturbations Reveal Synergy-Mediated, Context-Dependent Feedforward and Feedback Mechanisms of Adaptation. Sci Rep 2020. 10.1038/s41598-020-61231-8.

[49] Chvatal SA, Ting LH. Common muscle synergies for balance and walking. Front Comput Neurosci 2013;7:43259. 10.3389/FNCOM.2013.00048/BIBTEX.

[50] De Marchis C, Schmid M, Bibbo D, Bernabucci I, Conforto S. Inter-individual variability of forces and modular muscle coordination in cycling: A study on untrained subjects. Hum Mov Sci 2013;32:1480–94. 10.1016/J.HUMOV.2013.07.018.

[51] D’Avella A, Bizzi E. Shared and specific muscle synergies in natural motor behaviors. Proc Natl Acad Sci U S A 2005. 10.1073/pnas.0500199102.

[52] Pitto L, Kainz H, Falisse A, Wesseling M, Van Rossom S, Hoang H, et al. SimCP: A Simulation Platform to Predict Gait Performance Following Orthopedic Intervention in Children With Cerebral Palsy. Front Neurorobot 2019;13:463173. 10.3389/FNBOT.2019.00054/BIBTEX.

[53] Vandekerckhove I, Leuven KU. Predictive simulations of common gait features in children with Duchenne muscular dystrophy n.d. 10.1016/j.gaitpost.2023.07.257.

[54] Fregly BJ. A Conceptual Blueprint for Making Neuromusculoskeletal Models Clinically Useful. Applied Sciences 2021, Vol 11, Page 2037 2021;11:2037. 10.3390/APP11052037.

[55] Veerkamp K, van der Krogt MM, Waterval NFJ, Geijtenbeek T, Walsh HPJ, Harlaar J, et al. Predictive simulations identify potential neuromuscular contributors to idiopathic toe walking. Clin Biomech (Bristol, Avon) 2024;111. 10.1016/J.CLINBIOMECH.2023.106152.

[56] Van Wouwe T, Ting LH, De Groote F. An approximate stochastic optimal control framework to simulate nonlinear neuro-musculoskeletal models in the presence of noise. PLoS Comput Biol 2022;18:e1009338. 10.1371/JOURNAL.PCBI.1009338.

[57] Meyer AJ, Eskinazi I, Jackson JN, Rao A V., Patten C, Fregly BJ. Muscle synergies facilitate computational prediction of subject-specific walking motions. Front Bioeng Biotechnol 2016;4. 10.3389/fbioe.2016.00077.

[58] Veerkamp K, Carty CP, Waterval NFJ, Geijtenbeek T, Buizer AI, Lloyd DG, et al. Predicting Gait Patterns of Children With Spasticity by Simulating Hyperreflexia. J Appl Biomech 2023;39. 10.1123/jab.2023-0022.

[59] Hicks JL, Uchida TK, Seth A, Rajagopal A, Delp SL. Is My Model Good Enough? Best Practices for Verification and Validation of Musculoskeletal Models and Simulations of Movement. J Biomech Eng 2015;137. 10.1115/1.4029304/371243.

[60] Masel J. Genetic drift. Current Biology 2011;21:R837–8. 10.1016/J.CUB.2011.08.007.

[61] Tomasi M, Artoni A. Identification of Motor Control Objectives in Human Locomotion via Multi-Objective Inverse Optimal Control. J Comput Nonlinear Dyn 2023;18. 10.1115/1.4056588/1155848.

[62] Delp SL, Loan JP, Hoy MG, Zajac FE, Topp EL, Rosen JM. An Interactive Graphics-Based Model of the Lower Extremity to Study Orthopaedic Surgical Procedures. IEEE Trans Biomed Eng 1990;37:757–67. 10.1109/10.102791.

[63] Geijtenbeek T. SCONE: Open Source Software for Predictive Simulation of Biological Motion. J Open Source Softw 2019. 10.21105/joss.01421.

[64] Hyfydy - High Fidelity Dynamics n.d. https://hyfydy.com/ (accessed August 7, 2024).

[65] McCrea DA, Rybak IA. Organization of mammalian locomotor rhythm and pattern generation. Brain Res Rev 2008. 10.1016/j.brainresrev.2007.08.006.

[66] Kozlov AK, Lansner A, Grillner S, Kotaleski JH. A hemicord locomotor network of excitatory interneurons: A simulation study. Biol Cybern 2007;96:229–43. 10.1007/S00422-006-0132-2/METRICS.

[67] Grillner S, Halbertsma J, Nilsson J, Thorstensson A. The adaptation to speed in human locomotion. Brain Res 1979;165:177–82. 10.1016/0006-8993(79)90059-3.

[68] Matsuoka K. Sustained oscillations generated by mutually inhibiting neurons with adaptation. Biol Cybern 1985;52. 10.1007/BF00449593.

[69] Russo A Di, Stanev D, Armand S, Ijspeert A. Sensory modulation of gait characteristics in human locomotion: A neuromusculoskeletal modeling study. PLoS Comput Biol 2021;17. 10.1371/journal.pcbi.1008594.

[70] Uchida TK, Seth A, Pouya S, Dembia CL, Hicks JL, Delp SL. Simulating ideal assistive devices to reduce the metabolic cost of running. PLoS One 2016;11:1–19. 10.1371/journal.pone.0163417.

[71] Van Criekinge T, Saeys W, Truijen S, Vereeck L, Sloot LH, Hallemans A. A full-body motion capture gait dataset of 138 able-bodied adults across the life span and 50 stroke survivors. Scientific Data 2023 10:1 2023;10:1–8. 10.1038/s41597-023-02767-y.

[72] Lee D, Seung H. Algorithms for non-negative matrix factorization. Adv Neural Inf Process Syst 2001:556–62. 10.1109/IJCNN.2008.4634046.

